# Fruit flies actively restart their circadian clock by proactively shaping their environment

**DOI:** 10.1101/2025.03.26.645468

**Authors:** Angelica Coculla, Luis Garcia Rodriguez, Maite Ogueta, Ralf Stanewsky

## Abstract

Circadian clocks are prevalent on Earth and are generally believed to provide adaptive advantage to organisms. Functional circadian clocks, and their synchronization with the outside world, has also been implicated to provide health benefits for humans. However, experimental evidence for the benefits of possessing a circadian clock is sparse and largely restricted to prokaryotic organisms. Here, we provide evidence for the benefits of circadian clocks and temporally organized life in the fruit fly *Drosophila melanogaster*. We demonstrate that flies prefer and actively choose to live under circadian clock regulation: Exposure to constant light breaks down the circadian clock and leads to arrhythmic locomotor activity patterns. When given the choice to move between dark and illuminated areas in a constant light environment, flies were able to maintain, or even re-gain, rhythmic behavioural patterns. These rhythms were mirrored by regular positional changes between the two areas, demonstrating that flies actively contribute to creating an environment allowing circadian clock function and temporal organisation. The self-inflicted rhythms were accompanied by molecular rhythms in the majority of the clock neurons known to drive behavioural rhythms in flies, showing that they are indeed controlled by the circadian clock. Finally, behavioural rhythmicity was correlated with restoration of rhythmic sleep patterns and less-fragmented sleep compared to arrhythmic flies. While life is possible without a circadian clock, we show that if given the choice, animals prefer to live in a temporally organized manner and actively contribute to make this possible. This provides strong arguments for the benefits of possessing and using a circadian clock, for example by ensuring a better quality of sleep.

## Main Text

Circadian clocks are a universal feature of almost all organisms on our planet, driving daily rhythms of physiology and behaviour, even in the absence of rhythmic environmental cues of light and temperature. They independently evolved several times in life history due to the constant exposure to the diurnal changes of day and night (*1, 2*). Together with their omnipresence, this suggests that circadian clocks offer a substantial fitness advantage (‘adaptive value’), although this is rather difficult to demonstrate experimentally. Mutations abolishing circadian clock function are perfectly viable and fertile, and can be maintained in the laboratory without any problems (e.g., (*3*)). Although there are reports that clock mutants reduce lifespan and fecundity, these studies are complicated by the fact that most clock genes have known or potential pleiotropic functions in addition to their role in circadian timing (*4*–*6*). Convincing evidence for the fitness advantage of circadian clocks stems from growth competition experiments in Cyanobacteria, showing that period-altering clock mutants can outcompete wild type bacteria, as long as the period of the clock mutant matches that of the external light:dark (LD) cycle (*7, 8*). Likewise, mice with a fast-running clock were outcompeted by wild type mice after keeping them in several generations in a semi-natural habitat (*9*). However, these advantages are restricted to cyclic environmental conditions, and the adaptive value of possessing a circadian clock that operates in constant conditions remains ambiguous (*10*).

In most circadian assays measuring locomotor activity of rodents and insects, animals are exposed to a certain environmental condition (e.g., a standard 12 hr : 12 hr LD cycle), without giving them the opportunity to escape or to alter this environment. The animal can only adjust its behaviour to the specific condition, for example by being more active in the dark versus light condition for nocturnal animals. In these fixed environmental settings, *Drosophila melanogaster* exhibit crepuscular behaviour during LD cycles, and maintain rhythmic behaviour in constant darkness (DD) with a free-running period of approximately 24 hr, demonstrating that locomotor activity rhythms are controlled by a circadian clock (*3, 11*). In contrast, exposure of the flies to constant light (LL) leads to arrhythmic behaviour, due to the constant degradation of the clock protein Timeless (Tim), mediated by the blue-light photoreceptor Cryptochrome (Cry) and the F-Box protein Jetlag (Jet) (*12*–*14*). However, these classical assays do not reveal how animals behave in more natural conditions, in which they would be able to hide from bright light or warm temperatures, for example. To circumvent this problem, we designed a behavioural assay allowing flies to choose spending time in the light or dark. We wondered, if flies are able to actively restore their circadian clock and rhythmic behaviour, if they are given a choice to spend time in a dark area while housed in a constantly illuminated incubator. If true, this would indicate that circadian clock function and rhythmic behaviour are beneficial for the overall fitness of the fly, and that flies are actually able to perform proactive behaviours, i.e., that flies can actively change their behaviour to create a suitable environment, instead of only reacting to it.

## Results

We designed the choice apparatus based on the Ethoscope (*15*), a real-time and video-recording hardware that utilizes centroid analysis to quantify position and activity of flies (Fig 1A, 1B) (*12, 13*). The dark area was created by covering half of the arena with a high-pass infrared material to facilitate fly silhouette identification (Fig 1A). Flies were individually housed inside two glass tubes, with the open ends loosely connected to allow both air-exchange and passage of the fly between the tubes (Fig 1A, 1B). To prevent a light-independent preference for one tube, food was provided at both ends of the tubes (Fig 1B). Flies were first exposed to three full days of LD, after which each arena was subjected to one of the four different conditions: (A) flies were exposed to LL without being given the choice between light and dark (no cover). (B) flies were exposed to LL for three days before the cover was placed above one half of the arena, allowing them to choose between light and dark. (C) and (D) same as B, but the cover was placed above half of the arena immediately after the transition to LL (C), or already during the LD period (D) (Fig. 1C).

**Fig. 1.**
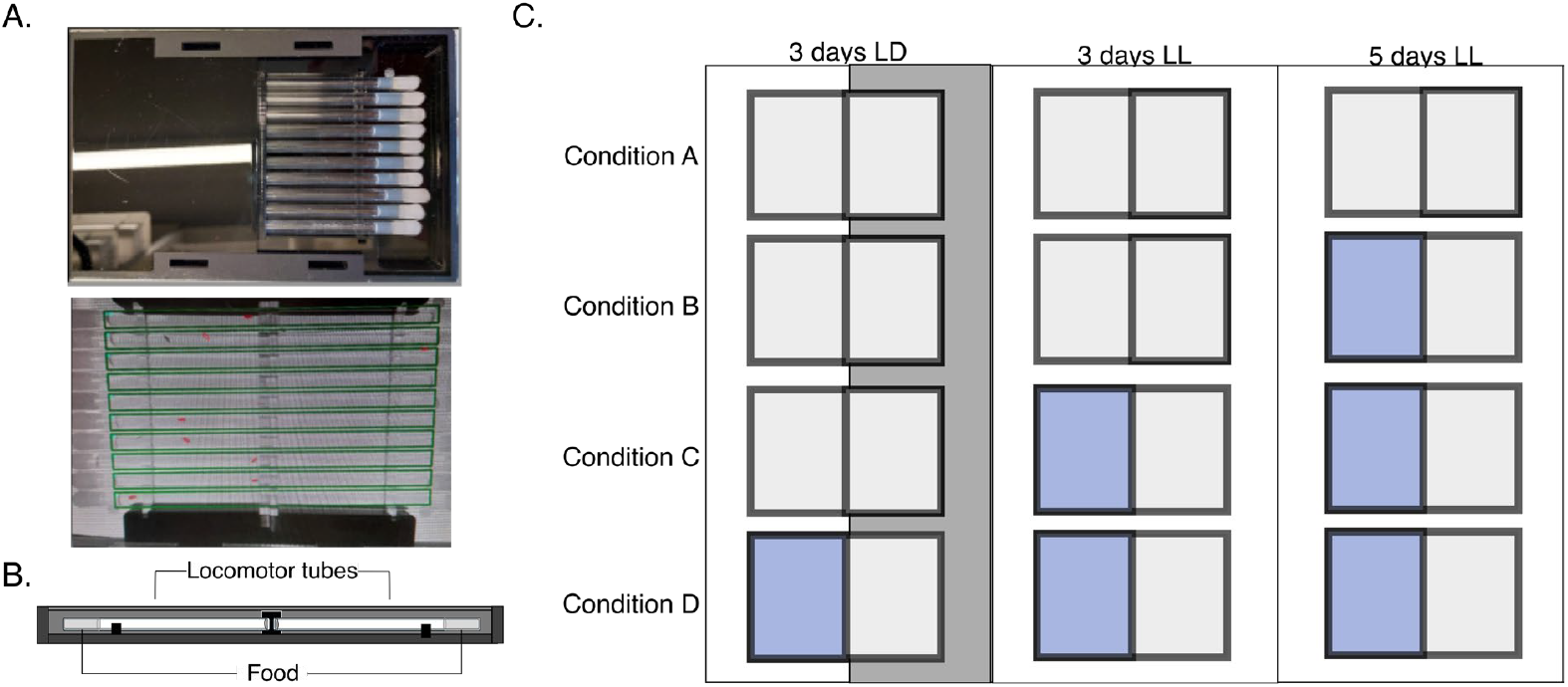
Experimental set-up of the light choice assay. (**A**) Eye (top) and camera (bottom) view of the ethoscope set-up. (**B**) Schematic of the locomotor tube assembly in the light choice assay. (**C**) Representation of the four conditions in the light-choice assay. The incubator light conditions are represented by the outer rectangles (light and dark periods are indicated by the white and grey rectangles, respectively). Covered areas (providing a dark sector) are represented by the blue square.

### Access to a dark area restores rhythmic behaviour in constant light

As expected, flies in all conditions showed crepuscular behaviour during the initial LD entrainment, with no preference for a particular side (Figs 2A, 3A, S1). Also as expected, almost all flies without a cover became arrhythmic during LL (Fig 2A, left, Tab 1) (*12, 13*). Strikingly, giving flies the opportunity to seek a dark area in LL restored or maintained robust circadian behaviour in ∼ 60% to 80% of individuals, depending on the condition, showing that flies that are given the choice between a dark and light area have a higher probability to be rhythmic than the flies without the cover (Fig 2A, 2B; Tab 1, S3) and with a stronger rhythm (RS > 0.1) than flies without the cover (Fig 2C; Tab S2, S4). Importantly, even in condition B, where flies initially become arrhythmic in LL, adding the cover after three days in LL restored rhythmicity in > 60 % of the flies, indicating that the flies actively chose spending time in the dark area, thereby restoring rhythmic circadian behaviour (Fig 2A-2C, Tab 1, S2, S4, S5). Independent of the condition, rhythmic flies in LL exhibit long free-running periods ranging between 27-29 hr (Fig 2B, 2D; Tab 1, S2, S4), which is substantially longer compared to the period in DD (Tab 1). The long rhythms are reminiscent of similarly long periods observed for wild type flies exposed to dim LL (1.5 lux) in classical, non-choice behavioural assays (*13*). This suggests that in our choice assay the flies enter the dark area of the arena only to a certain extent, still exposing them to dim light—enough to cause a period lengthening, but insufficient to cause arrhythmicity (see below).

**Fig. 2.**
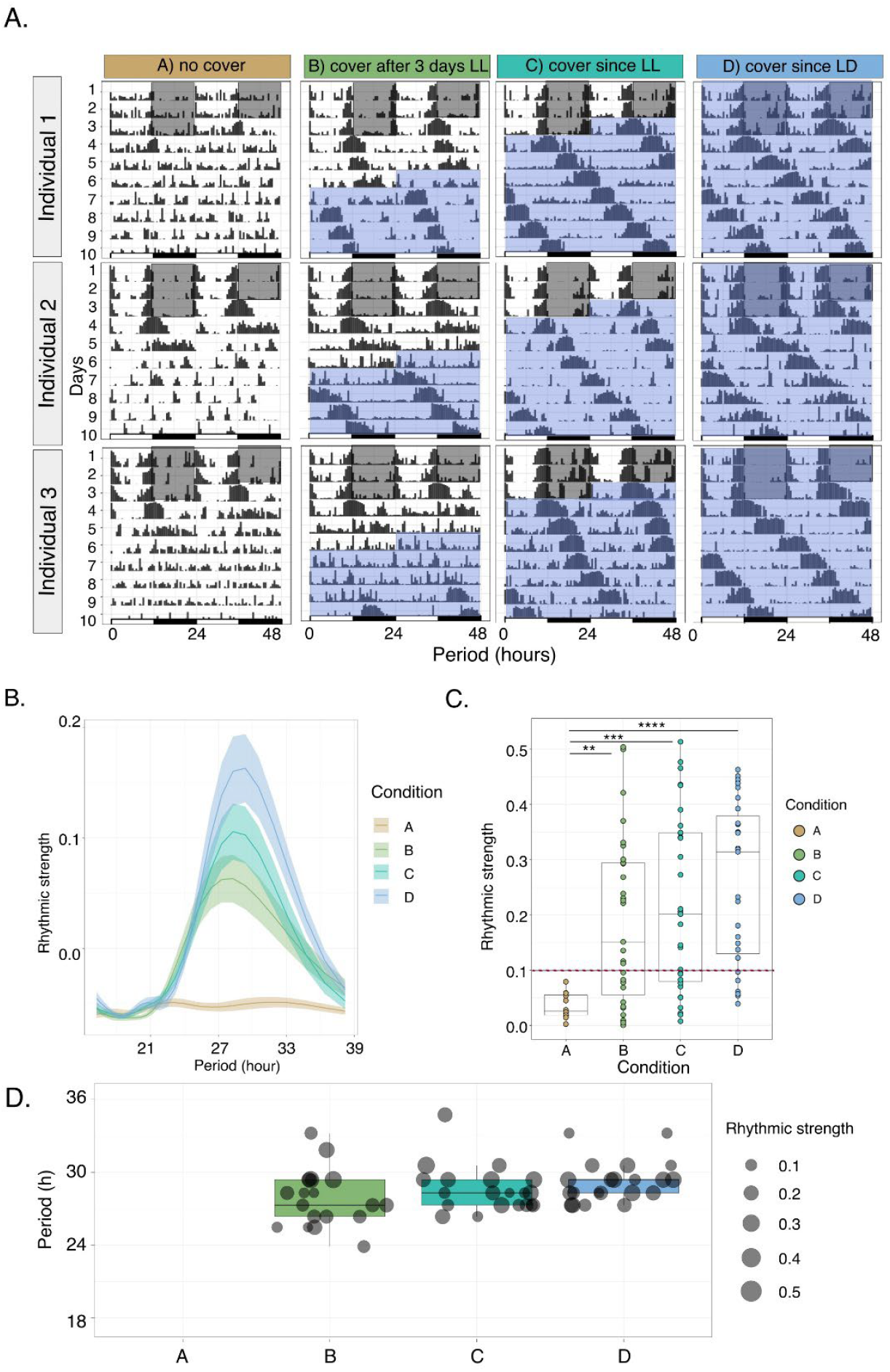
Flies with a choice maintain or re-establish rhythmic behavior in LL. (A) Representative individual actograms for each condition. Blue areas indicate the days where the cover was added to one side of the arena. (**B**) Population periodogram for each condition from LL4 to LL8 (GLM, family: binomial, *p<* .0001; Tab S3). (**C**) Rhythmic strength (Lomb Scargle, *alpha =* .01) of the rhythmic flies. The red dashed line represents the cut-off to classify rhythms in strong and weak (Levene’s test: *F =* 7.77, *p <* .001; Welch ANOVA: *F*_(3,51.51)_ = 38.12, *p <* .0001; Tab S2, S4; Games-Howell test see Tab S5). (**D**) Period length of flies with a strong rhythm in each condition (Levene’s test: *F* = 6.72, *p <* .001; Welch ANOVA: *F*_(3,37.03)_ = 2.51, *p* > .05; Tab S2, S4; Games-Howell test see Tab S5). **** p < .0001, *** p < .001, ** p < .01, * p < .05, n.s. p >.05.

**Table 1.** Rhythmicity, period length, and rhythmic strength of flies in LL and DD.

| *Rhythmicity after 4 days in constant light (LL)* |  |  |  |
| --- | --- | --- | --- |
| Condition | % rhythmic flies (# rhythmic flies/total) | Period (h) ± SEM | Power ± SEM |
| No cover (A) | 28% (12/43) | 25.8 ± 1.73 | 0.1 ± 0.06 |
| Cover after 3 days LL (B) | 62% (31/50) | 27.1 ± 0.60 | 0.3 ± 0.03 |
| Cover since LL (C) | 67% (31/46) | 28.1 ± 0.42 | 0.2 ± 0.03 |
| Cover since LD (D) | 77% (27/35) | 28.9 ± 0.49 | 0.3 ± 0.03 |
| *Rhythmicity in constant darkness (DD)* |  |  |  |
| Constant darkness (DD) | 100% (15/15) | 24.3 ± 0.11 | 0.3 ± 0.02 |

Rhythmicity, period and power were assessed using the Lomb-Scargle algorithm ( $\alpha=.01$ ).

### Flies rhythmically enter the dark area when exposed to constant light

The results described above indicate that flies exposed to LL indeed use the opportunity to enter the dark area for restoring rhythmicity in LL. To test how often and to which extend they enter the dark area, we determined the localization of each fly during the entire experiment for each of the four conditions (Fig 3A). As expected, during the initial LD period flies did not show any clear preference for either side of the tube, independent of the presence of a cover. However, flies exhibiting rhythmic behaviour in LL due to the presence of a cover, showed a preference for the dark area, while rhythmically approaching, and occasionally entering, the light area (Fig 3A). Interestingly, flies behaving arrhythmically in the same condition, did not show this spatial preference rhythm, demonstrating that rhythmic relocation between both areas is causally linked to rhythmic locomotor activity (Fig. 3A). Analysing the period length of the spatial distribution rhythm, further supports the interdependency with the locomotor activity rhythms, since both have a similarly long period (Fig 2B, 3B) and a stronger rhythmic strength (Fig 2C, 3C; Tab S2, S4, S5) in all conditions inducing rhythmicity. To confirm this, we compared period length and rhythmic strength of locomotor activity and position change rhythms of all rhythmic flies (Fig 4A-4D). Indeed, period length and rhythmic strength of both rhythms were not significantly different in all conditions (Fig 4A, 4C, Tab S2, S4). Similarly, we observed a strong correlation in the rhythm strength between activity and position rhythms (Fig 4D, Tab S4), but only a weak correlation for period length (Fig 4B, Tab S4). These results emphasize the importance of rhythmicity *per se*, rather than precise measurement of period length.

**Fig. 3.**
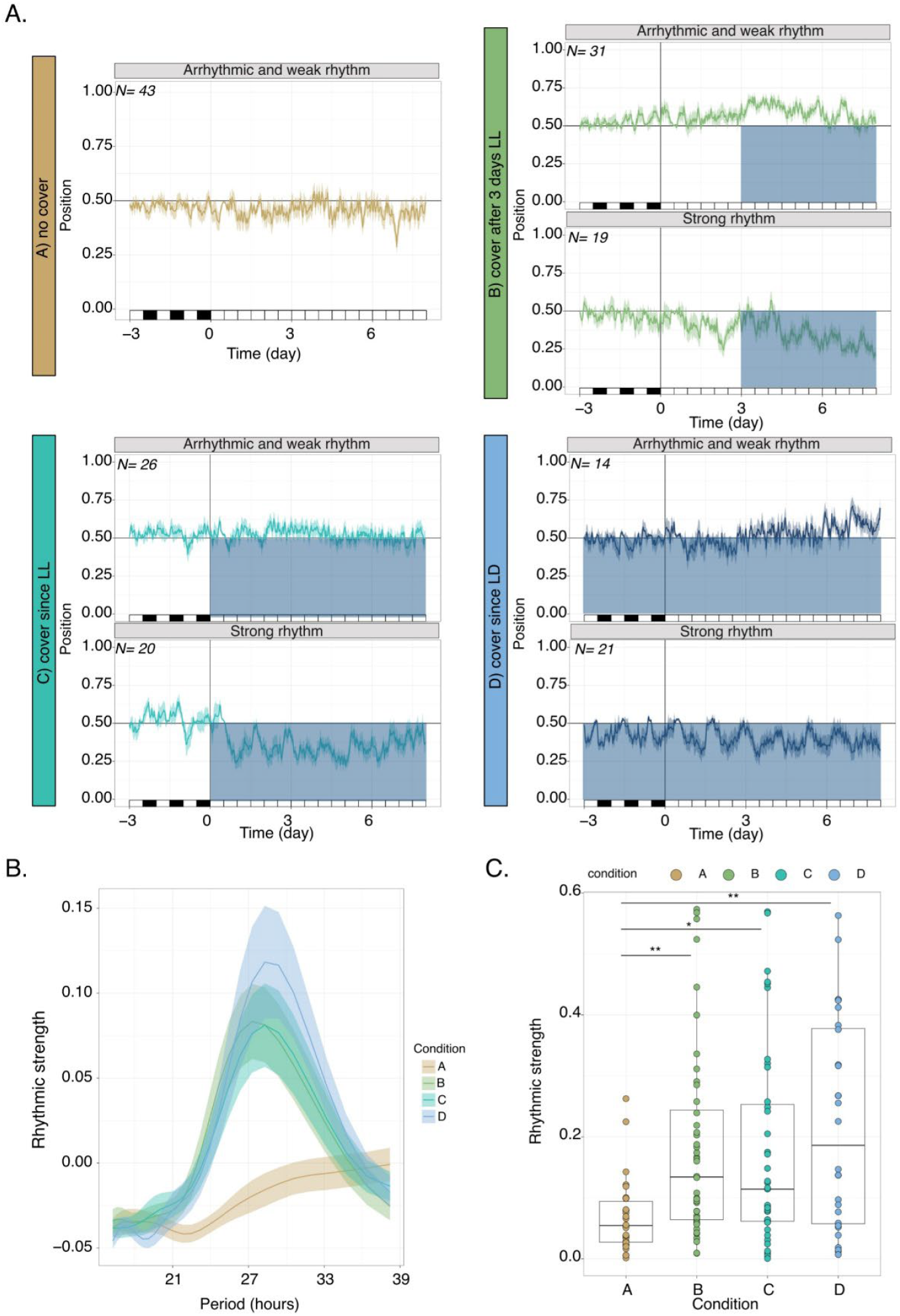
Rhythmic flies approach the illuminated area in a rhythmic manner. **A**) Overall position of the flies across experimental days in arrhythmic and rhythmic flies for each condition. Blue areas indicate the covered area of the arena. (**B**) Average period length of flies in each condition (GLM, family = binomial, *p >* .05; Tab S3). (**C**) Rhythmic strength of positional change within the tube across conditions (Levene’s test: *F* = 6.26, *p <* .001; Welch ANOVA: *F*_*(3,68*.*37*)_= 13.08, *p* > .0001; Tab S2, S4; Games- Howell test see Tab S5). ** p < .01, * p < .05, n.s. p >.05.

**Fig. 4.**
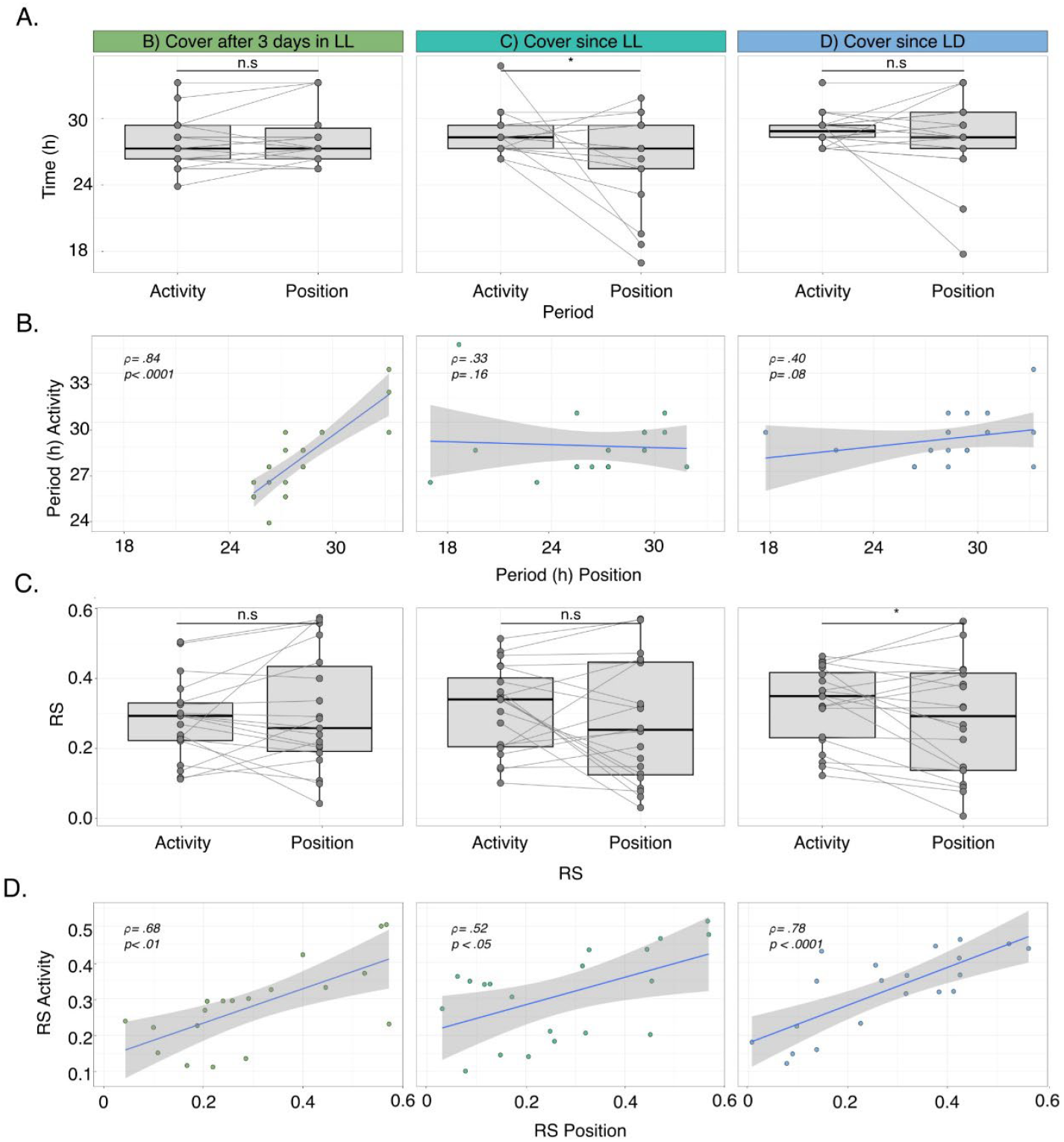
Strength of locomotor rhythmicity is correlated with rhythmic strength of positional change. (**A**) Boxplots showing differences in period length of locomotor activity and position change for individual flies, connected by grey lines (Levene’s test: *F* = 17.32, *p <* .0001; paired *t-*test with Welch correction: Condition B: *t(37)=* -1.83, *p* >.05; Condition C: *t(39)=* 2.85, *p* <0.05; Condition D: *t(39)=* 1.11, *p* >.05; Tab S2, S4). (**B**) Correlation between locomotor activity and position change of each fly based on period length (Tab S4). (**C**) Boxplots showing differences in rhythmic strength of locomotor activity and position change for individual flies, connected by grey lines (Levene’s test: *F* = 16.16, *p <* .0001; paired *t-*test with Welch correction: Condition B: *t(37)=* -1.03, *p* >.05; Condition C: *t(39)=* 1.64, *p* >.05; Condition D: *t(39)=* 2.67, *p* <.05; Tab S2, S4). (**D**). Correlation between locomotor activity and position change of each fly based on period length. Correlation coefficients (ρ) and p-values are shown for each condition (Tab S3). Only flies with a strong rhythm were included for clarity reasons (condition B: N=19; condition C: N=20, condition D: N=21), but inclusion of all rhythmic individuals produced similar results (not shown). ** p < .01, * p < .05, n.s. p >.05.

### Flies able to generate rhythmic behaviour in constant light exhibit Period oscillations in clock neurons

If the rhythmic LL-behaviour observable in the majority of flies with access to a dark area is indeed due to reinitiated or maintained clock function, molecular oscillations of clock gene products should occur in the clock neurons of these flies. Because rhythmic flies are not synchronized to each other and exhibit variably long behavioural periods (Fig 2A, Tab 1) it is not straight-forward to predict peak and trough phases of molecular oscillations for individual flies, which is necessary to reveal potential molecular oscillations. To circumvent this problem, we observed the behaviour of individual flies kept in condition C up to day 4 in LL, and after extrapolating their activity pattern or the next day, we sacrificed flies at times they were predicted to be active or inactive on day 5 in LL. We chose to dissect the flies on day 5 in LL, to rule out that dampening Period oscillations, originally synchronized to the initial LD phase, contribute to any oscillations observed in LL (PER oscillations cease after 2 days in LL: (*14*)). We then compared PER levels across all clock neuronal groups between flies predicted to be in the middle of their active or inactive phase, respectively. For all neuronal groups we found higher PER levels in the inactive flies compared to the active flies (Fig 5A, 5B, S2). We used Estimation Statistics (ES), which focusses on the magnitude of the effect size (i.e., the mean difference) and its precision, and therefore offers a more informative way to interpret results compared to tests restricted to acceptance or rejection of the null-hypothesis to pairwise compare staining intensity between active and inactive flies (*16*) (Fig 5A). The top part of the Multi two-group Cumming plot shows the individual data points, with the mean and standard error plotted to the right (the gap indicates the mean), while the lower part shows the mean difference (black dots), the resampled distribution (coloured curves), as well as the bootstrap 95% confidence interval (CI, vertical black lines). The CI of the LNd, the l-LNv, and the DN3, are above the reference line, while in the s-LNv and DN2 it marginally crosses the line (−0.17 and -0.37, respectively; Tab S6) indicating a difference between PER levels between inactive and active flies in these neurons (Fig 5A, S2; Tab S6). In the other neuronal groups (5^th^ s-LNv, DN1), there is also a trend for higher PER intensity in the inactive flies, but the difference is below the 95% CI (Fig 5A, S2; Tab S6). These results fit well with data reported for flies kept in DD, where peak PER expression occurs in the transition from the subjective night to the subjective day, when flies are inactive, while at the end of the subjective day, when flies are active, PER levels are low (*17*–*19*).

**Fig. 5.**
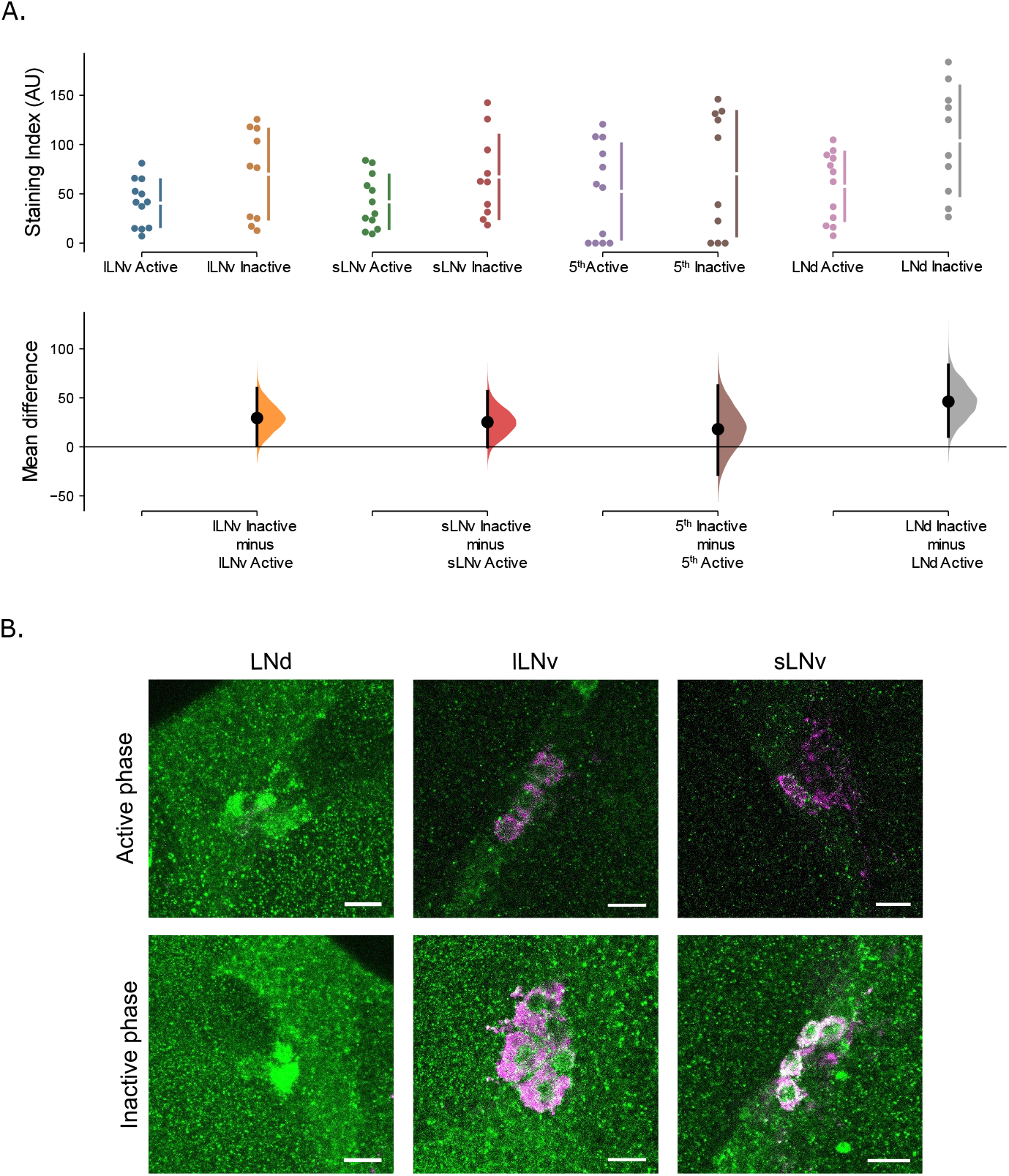
Flies maintaining rhythmicity in LL display PERIOD oscillations in clock neurons. (**A**) Quantification of PER expression in the different lateral neuronal groups for rhythmic flies in LL (condition C), selected when they are either in their active or the inactive phase. The upper plot represents all the datapoints, with the mean (gap) and the standard deviation (length of vertical bars) for each neuronal group to the right. The lower plot shows the mean difference of the data from the inactive phase compared to the active phase as a bootstrap 95% confidence interval (see results text for details). (**B**) Representative images of the different groups of lateral neurons. Anti-PER: green, anti-PDF: magenta. Scale bar = 10μm. n number: active phase 12, inactive phase 10, from 3 independent runs.

### The ability to generate rhythmic behaviour in constant light correlates with less-fragmented sleep

A benefit from possessing a functional circadian clock is the ability to maintain a regular sleep pattern, which contributes to the overall fitness of organisms. To investigate if flies that actively maintain or restore circadian rhythmicity by regularly switching between light and dark portions of the locomotor tube also show a regular sleep pattern, we quantified and compared sleep parameters of flies displaying rhythmic or arrhythmic locomotor behaviour, kept in the same environmental condition (B-D, see Fig 1). First, to rule out any sleep altering factors in the genetic background or in our experimental set-up, we analysed sleep for the LD part of the experiment and found the typical day and night sleep patterns in all four conditions (Fig S3). As expected, in the LL part of the experiment rhythmic flies also showed a circadian sleep pattern (Fig 6A). In contrast, arrhythmic or weakly rhythmic flies (including flies kept in condition A), showed no daily variation in their sleep pattern, although total sleep amount was not affected (Fig 6A, S3, Tab S1). To determine sleep quality, we also analysed the number of sleep bouts, their average duration, and the duration of the longest sleep bout during days 4-8 after transfer to LL. Rhythmic flies kept in condition D showed a clear improvement of sleep quality, indicated by a significant decrease in the total number of sleep bouts and significant increases of the average and longest sleep bout duration compared to the arrhythmic flies (Fig 6B, Tab S1, S2, S4). The results for rhythmic flies kept in condition C were similar, although the average duration of sleep bouts was not significantly different between rhythmic and arrhythmic flies (Fig. 6B, Tab S1, S2, S4). Importantly, the median duration of the longest sleep bout was doubled in rhythmic versus arrhythmic flies kept in condition C (∼4 h vs ∼2 h, respectively, Fig 6B, Tab S1). Flies that restarted rhythmic behaviour (condition B) did not show a significant improvement of sleep quality compared to the arrhythmic flies in this condition, however—as in condition C and D—the longest sleep bouts were observed in rhythmic flies (Fig. 6B, Tab S1, S2, S4). Moreover, compared to the arrhythmic flies in condition A, flies that restarted rhythmic locomotor behaviour (condition B) also exhibited a robust circadian sleep pattern (Fig. 6A, S3). Taken together, the sleep analysis shows that flies that maintained rhythmic behaviour in LL by actively relocating between the illuminated and dark areas of the arena, display higher quality (i.e., less fragmented) sleep compared to their arrhythmic counterparts. Flies in condition B, despite developing a circadian sleep pattern after providing them with access to a dark area, presumably need more time to improve their sleep quality, indicating that rhythmic locomotor behaviour and sleep, does not immediately translate to higher quality sleep.

**Fig. 6.**
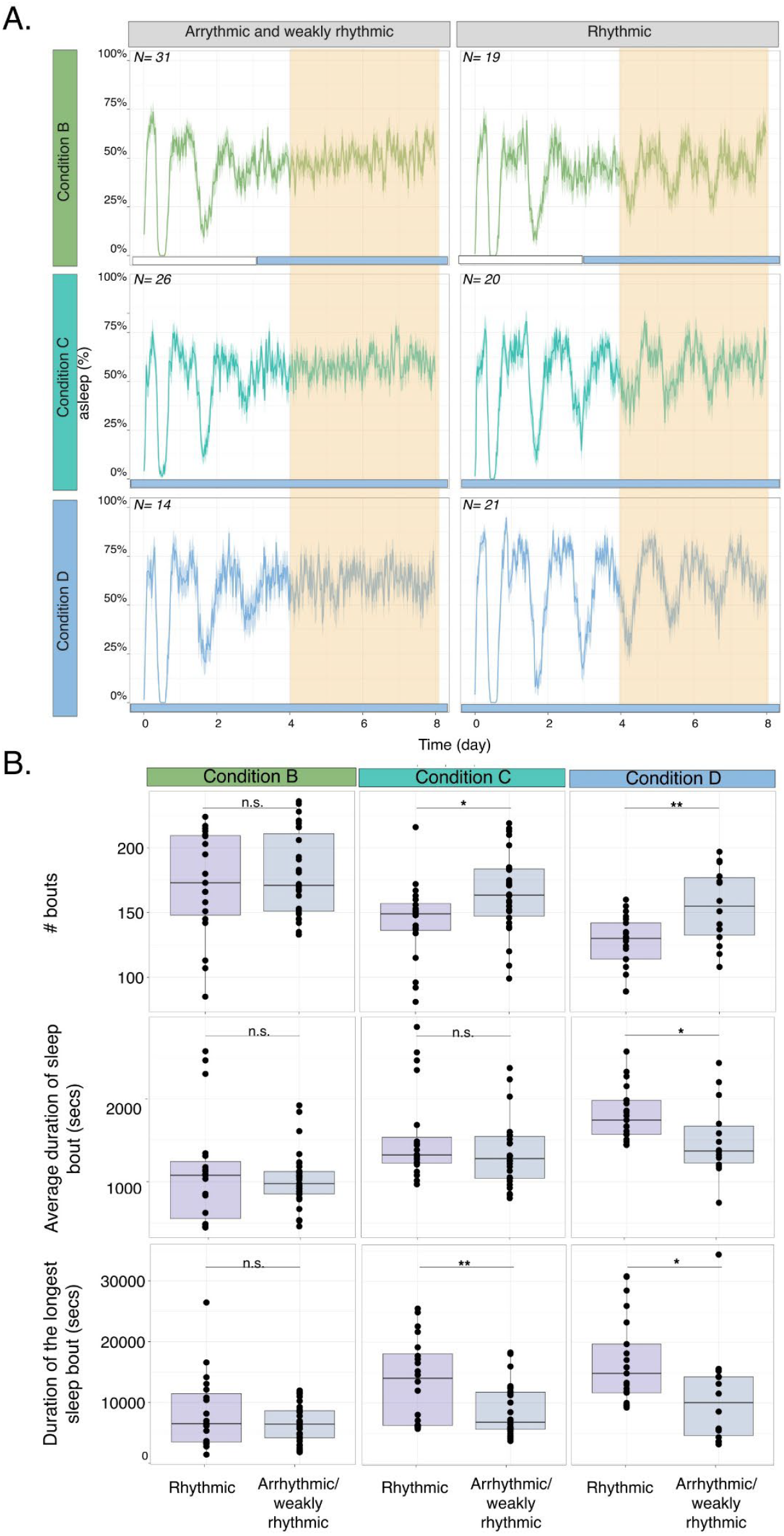
Flies maintaining rhythmic locomotor activity in LL have less fragmented sleep. (**A**) Population sleep patterns in LL. Yellow areas indicate days during which sleep was analysed (days 4 to 8 in LL). White and blue bars below the sleep plots indicate days when the cover was applied. (**B**) Comparison of sleep parameters between flies restarting or maintaining rhythmic locomotor activity in LL and arrhythmic or weakly rhythmic flies (Tab S2, S4). ** p < .01, * p < .05, n.s. p >.05.

## Discussion

We show here that *Drosophila melanogaster* surprisingly are capable of performing proactive behaviours. Arrhythmic flies avoid arrythmia-inducing LL exposure, by periodically visiting the dark area of the behavioural test tube, thereby ensuring molecular circadian clock function as well as rhythmic locomotor activity and sleep. Flies do not need a functional circadian clock or rhythmic locomotor behaviour in order to survive or reproduce (*3*). Therefore, it is quite surprising to see that, if given the choice, flies ‘decide’ to restart their clock, allowing them to perform circadian locomotor behaviour. We think that this behaviour demonstrates the importance of circadian clock function. Our results show that self-established circadian rhythmicity improves the quality of sleep, which could be a subsequent benefit of regained rhythmicity. Metabolism in insects and mammals is also under circadian control (*20, 21*). Therefore, the metabolism of the rhythmic flies is probably also temporally organized and operates more economically compared to the arrhythmic flies, providing a further benefit.

But do the flies in condition B (Fig 2A) indeed ‘restart’ their circadian clock upon given the choice to enter the dark area? This is a bit of a chicken and egg problem: How can an arrhythmic fly rhythmically enter the dark or illuminated area to start its clock? It is possible that one, perhaps random visit to the dark area is sufficient to restart the molecular clock, which could then lead to rhythmic behaviour. It is known that dark pulses can synchronize circadian clocks in mammals and plants (*22, 23*). *Drosophila melanogaster* does prefer dim light (*24*), making this scenario possible. Alternatively, in some individuals there might be some remaining clock function present before the cover is placed on the arena, allowing these individuals to rhythmically switch between the dark and illuminated area and thereby to re-establish strong circadian oscillations. The fact that we observed a higher degree of rhythmicity in the conditions where the cover was placed on the arena earlier, before exposing the flies to LL (conditions C, D, Fig 2A, Tab 1), argues that the stronger rhythms in these flies helped them to rhythmically visit the dark area in LL. In any case, the results show that flies proactively change their behaviour in order to maintain (conditions C, D) or even restart (condition B) circadian behaviour, by rhythmically switching between the dark and illuminated areas and thereby creating their own LD cycle.

Flies able to restart their locomotor-rhythm after being arrhythmic (condition B) also showed rhythmic sleep patterns (Fig 6A, S3), but without showing an improved quality of sleep (Fig 6B). We think this shows that rhythmic locomotor activity and sleep not immediately translates to a better quality of sleep. Flies that do show an improvement of sleep (condition C and D) were already rhythmic for the entire LD and LL part of the experiment (6 days) before their sleep was assessed (days 4-6 in LL), while the condition B flies had just restarted their rhythms in this period (Fig 6A). It is therefore possible that a prolonged time of rhythmic behaviour is required to develop a less-fragmented, higher quality sleep. It would be interesting to determine if the quality of sleep in mammals is also linked to the history of previous rhythmic behaviour. If true, this would emphasize the importance of clock synchronization and regular daily sleep routines for sleep quality in humans, and add an important factor to the growing list of health risks associated with rotating shift work (*25, 26*).

Depending on the condition, between 60% - 80% of the flies were able to behave rhythmically in LL conditions by generating their own LD cycle (Fig. 2B). What distinguishes these flies from those that behave arrhythmic in LL? One possibility is that rhythmic individuals share a genetic trait enabling them to maintain or re-establish rhythmic behaviour under LL choice conditions. Even though the chromosomes of wild type stock we used (*iso31*) have been isogenized, resulting in limited genetic variation, it is still possible that genetic differences exist, leading to the observed behavioural variation. It is also possible that individual variation has no genetic component, as described for walking behaviour patterns in *Drosophila melanogaster* (*27*). The authors show in this study that stochastic morphological differences in the wiring of visual system neurons leads to the individualized behavioural patterns. In both scenarios, retesting the same individuals in the same choice assay should give the same result, i.e., flies behaving rhythmic should again behave rhythmic and arrhythmic flies should again stay arrhythmic. Alternatively, minimal differences in the local environmental conditions (light-intensity, shade, etc) could contribute to the observed differences, although we did not observe any reproducible correlations between position in the incubator and rhythmicity. Recently it has been shown that populations of the red flour beetle *Tribolium castaneum* show a similar degree of variability in rhythmic behaviour, both in DD and LL (*28*). It is therefore indeed possible, that either genetic, or stochastic differences between individuals contribute to behavioural variability by tipping the balance toward clock-controlled, or arrhythmic behaviour in certain conditions. Such individual variability could offer an adaptive value for a given individual under a certain environmental condition, thereby contributing to the overall flexibility and adaptation of the population (*29*).

(*30*) (*31*) (*32*) (*33*)

## Acknowledgments

### Funding

This work was funded by the German Research Foundation (DFG) as part of the SFB TRR 212 (NC3, Project number 316099922)—Project number 471672756 (to Ralf Stanewsky).

## Author contributions

A.C. and R.S. designed the study. A.C. performed experiments and analyzed the data. L.G. designed and built the behavioral choice apparatus. M.O. performed the immunostainings and quantification of the images. A.C. and R.S. wrote the manuscript.

## Competing interests

The authors have no competing interests.

## Data and materials availability

All original raw data files are available from the corresponding author upon request.

## Supplementary Materials

Materials and Methods

Figs. S1 to S4

Tables S1 to S6

## Notes

### Competing Interest Statement

The authors have declared no competing interest.

## References and Notes

1. W. Gehring, M. Rosbash, The Coevolution of Blue-Light Photoreception and Circadian Rhythms. Journal of Molecular Evolution 57, S286–S289 (2003).

2. U. Bhadra, N. Thakkar, P. Das, M. Pal Bhadra, Evolution of circadian rhythms: from bacteria to human. Sleep Medicine 35, 49–61 (2017).

3. R. J. Konopka, S. Benzer, Clock mutants of Drosophila melanogaster. Proc Natl Acad Sci U S A 68, 2112–6 (1971).

4. L. Beaver, B. Gvakharia, T. Vollintine, D. Hege, R. Stanewsky, J. Giebultowicz, Loss of circadian clock function decreases reproductive fitness in males of Drosophila melanogaster. Proc National Acad Sci 99, 2134–2139 (2002).

5. M. Horn, O. Mitesser, T. Hovestadt, T. Yoshii, D. Rieger, C. Helfrich-Förster, The Circadian Clock Improves Fitness in the Fruit Fly, Drosophila melanogaster. Front Physiol 10, 1374 (2019).

6. A. Klarsfeld, F. Rouyer, Effects of circadian mutations and LD periodicity on the life span of Drosophila melanogaster. Journal of biological rhythms 13, 471–478 (1998).

7. Y. Ouyang, C. Andersson, T. Kondo, S. Golden, C. Johnson, Resonating circadian clocks enhance fitness in cyanobacteria. Proc Natl Acad Sci U S A 95, 8660–8664 (1998).

8. M. A. Woelfle, Y. Ouyang, K. Phanvijhitsiri, C. H. Johnson, The adaptive value of circadian clocks: an experimental assessment in cyanobacteria. Curr Biol 14, 1481–6 (2004).

9. K. Spoelstra, M. Wikelski, S. Daan, A. S. I. Loudon, M. Hau, Natural selection against a circadian clock gene mutation in mice. Proc Natl Acad Sci U S A 113, 686–691 (2016).

10. M. L. Jabbur, C. Dani, K. Spoelstra, A. N. Dodd, C. H. Johnson, Evaluating the Adaptive Fitness of Circadian Clocks and their Evolution. J Biol Rhythms 39, 115–134 (2024).

11. M. J. Hamblen-Coyle, D. A. Wheeler, J. E. Rutila, M. Rosbash, J. C. Hall, Behavior of period-altered circadian rhythm mutants of Drosophila in light:dark cycles. J. Insect Behav. 5, 417–446 (1992).

12. P. Emery, R. Stanewsky, J. Hall, M. Rosbash, A unique circadian-rhythm photoreceptor. Nature 404, 456–457 (2000).

13. R. Konopka, C. Pittendrigh, D. Orr, Reciprocal behaviour associated with altered homeostasis and photosensitivity ofDrosophilaclock mutants. J Neurogenet 6, 1–10 (1989).

14. N. Peschel, K. F. Chen, G. Szabo, R. Stanewsky, Light-dependent interactions between the Drosophila circadian clock factors cryptochrome, jetlag, and timeless. Curr. Biol. 19, 241–7 (2009).

15. Q. Geissmann, L. Garcia Rodriguez, E. J. Beckwith, A. S. French, A. R. Jamasb, G. F. Gilestro, Ethoscopes: An open platform for high-throughput ethomics. PLoS Biol 15, e2003026 (2017).

16. J. Ho, T. Tumkaya, S. Aryal, H. Choi, A. Claridge-Chang, Moving beyond P values: data analysis with estimation graphics. Nat Methods 16, 565–566 (2019).

17. C. Helfrich-Förster, Differential control of morning and evening components in the activity rhythm of Drosophila melanogaster--sex-specific differences suggest a different quality of activity. J Biol Rhythms 15, 135–54 (2000).

18. Y. Lin, G. D. Stormo, P. H. Taghert, The neuropeptide pigment-dispersing factor coordinates pacemaker interactions in the Drosophila circadian system. J Neurosci 24, 7951–7957 (2004).

19. O. T. Shafer, M. Rosbash, J. W. Truman, Sequential nuclear accumulation of the clock proteins Period and Timeless in the pacemaker neurons of Drosophila melanogaster. J Neurosci 22, 5946–54 (2002).

20. J. Bass, Circadian topology of metabolism. Nature 491, 348–356 (2012).

21. A. Sehgal, “Control of Metabolism by Central and Peripheral Clocks in Drosophila” in A Time for Metabolism and Hormones, P. Sassone-Corsi, Y. Christen, Eds. (Springer International Publishing, Cham, 2016; http://link.springer.com/10.1007/978-3-319-27069-2_4) Research and Perspectives in Endocrine Interactions, pp. 33–40.

22. H. Fukuda, H. Murase, I. T. Tokuda, Controlling Circadian Rhythms by Dark-Pulse Perturbations in Arabidopsis thaliana. Sci Rep 3, 1533 (2013).

23. J. Y. Mendoza, H. Dardente, C. Escobar, P. Pevet, E. Challet, Dark pulse resetting of the suprachiasmatic clock in Syrian hamsters: behavioral phase-shifts and clock gene expression. Neuroscience 127, 529–537 (2004).

24. W. Bachleitner, L. Kempinger, C. Wülbeck, D. Rieger, C. Helfrich-Förster, Moonlight shifts the endogenous clock of Drosophila melanogaster. Proc Natl Acad Sci U S A 104, 3538–43 (2007).

25. D. A. Bechtold, J. E. Gibbs, A. S. Loudon, Circadian dysfunction in disease. Trends Pharmacol Sci 31, 191–8 (2010).

26. D. B. Boivin, P. Boudreau, A. Kosmadopoulos, Disturbance of the Circadian System in Shift Work and Its Health Impact. J Biol Rhythms 37, 3–28 (2022).

27. G. A. Linneweber, M. Andriatsilavo, S. B. Dutta, M. Bengochea, L. Hellbruegge, G. Liu, R. K. Ejsmont, A. D. Straw, M. Wernet, P. R. Hiesinger, B. A. Hassan, A neurodevelopmental origin of behavioral individuality in the Drosophila visual system. Science 367, 1112–1119 (2020).

28. R. R. T. Prüser, N. K. E. Schulz, P. M. F. Mayer, M. Ogueta, R. Stanewsky, J. Kurtz, Deciphering a Beetle Clock: Individual and Sex-Dependent Variation in Daily Activity Patterns. J Biol Rhythms 39, 484–501 (2024).

29. I. A. Krams, T. Krama, R. Krams, G. Trakimas, S. Popovs, P. Jõers, M. Munkevics, D. Elferts, M. J. Rantala, J. Makņa, B. L. de Bivort, Serotoninergic Modulation of Phototactic Variability Underpins a Bet-Hedging Strategy in Drosophila melanogaster. Front Behav Neurosci 15, 659331 (2021).

30. K. Koh, W. Joiner, M. Wu, Z. Yue, C. Smith, A. Sehgal, Identification of SLEEPLESS, a sleep-promoting factor. Science 321, 372–376 (2008).

31. Q. Geissmann, L. Garcia Rodriguez, E. J. Beckwith, G. F. Gilestro, Rethomics: An R framework to analyse high-throughput behavioural data. PLoS One 14, e0209331 (2019).

32. R. Stanewsky, B. Frisch, C. Brandes, H.-C. Mj, M. Rosbash, J. Hall, Temporal and spatial expression patterns of transgenes containing increasing amounts of theDrosophilaclock geneperiodand alacZreporter: mapping elements of the PER protein involved in circadian cycling. The Journal of neuroscience : the official journal of the Society for Neuroscience 17, 676–696 (1997).

33. J. Schindelin, I. Arganda-Carreras, E. Frise, V. Kaynig, M. Longair, T. Pietzsch, S. Preibisch, C. Rueden, S. Saalfeld, B. Schmid, J.-Y. Tinevez, D. J. White, V. Hartenstein, K. Eliceiri, P. Tomancak, A. Cardona, Fiji: an open-source platform for biological-image analysis. Nature Methods 9, 676–682 (2012).

34. RStudio Team (2020). RStudio: Integrated Development for R. RStudio, PBC, Boston, MA URL http://www.rstudio.com/.

