## Supplementary Information for "Fruit flies actively restart their circadian clock by proactively shaping their environment"

### Materials and Methods

*Fly strains and maintenance.*

For all experiments we used the isogenized *Drosophila melanogaster* strain *w; iso31 (30)*. Flies were reared in plastic vials with food consisting of 0.7% agar, 1.0% soy flour, 8.0% cornmeal, 1.8% yeast, 8.0% malt extract, 4.0% sugar beet sirup, 0.8% propionic acid, 2.3% nipagen under 12 h:12 h LD cycles at 60% relative humidity and 25°C.

*Behavioral assay and hardware.*

Locomotor activity and position were recorded using modified Ethoscopes *(15)*. The light-dark choice apparatus consists of a flat plate (155 x 100 mm) that accommodates 10 pairs of standard locomotor tubes (~ 65 mm in length each) facing each other with their open ends, with *ad libitum* food (4% sucrose and 2% agar) located at the other end of each tube. One fly is loaded into each pair and tubes are loosely connected to prevent the fly from escaping, while allowing air exchange (Fig 1A). The behavioral apparatus and the cover used to create the dark area were built in house using a laser-cutter and high-pass infrared material (thickness: 3 mm, LUXACRYL-IR©, TTV GmbH). To facilitate the recording and to provide homogenous background infrared illumination, the assay was arranged on a custom-made light guide panel. It consists of a sandwich construction of transparent acryl (dimensions: 10 mm), infrared diffusor material (dimensions: 1 mm, LUXACRYL-STR3, opal 1175) and a reflective base made out of aluminum tape. The infrared light is generated by two stripes if IR LEDs (SOLAROX© LED STREIFEN INFRAROT 850NM IR1-60-850) along each side long side of the panel. Ethoscopes (maximum of four Ethoscopes in one I-30 PERCIVAL incubator) were evenly distributed along the plate to ensure equal light exposure. The detailed design of the behavioral arena is available upon request. One to four days old male flies were first entrained for at least three full days to standard 12hr:12hr light-dark cycles (LD), and then released into constant dim light (LL) (~ 4,5 -5,0 μW/m2, absolute spectral irradiance: 491 nm; ~ 20 lux) for a minimum of 8 days. We used two fluorescent light bulbs (17 Watt, F17T8ITL841, Alto II™ TECHNOLOGY, PHILIPS) located one at the back and one at the front of the incubator in opposite corners from one another. Light intensity was measured with an Ocean Optics QE65000by spectrometer, located in the center of the IR-diffuser plate between the four Ethoscopes. All flies were kept at a constant temperature (25°C) for the duration of the experiment. Experiments were repeated five times for all conditions, except for condition D (repeats: 4), because of a software crash during the recording. Data in constant darkness were recorded with the Drosophila Activity Monitor (DAM) system using DAM2 monitors (Trikinetics).

*Behavioral analysis.*

To determine period length and rhythmicity of activity recorded with the DAM system, data were extracted with a 30 minutes interval while Ethoscope data were extracted at 10 seconds intervals. In order to avoid potential effects of the different light transition (LD to LL, no cover to cover) and to allow comparisons between different conditions, we analyzed activity, position and sleep after day 4 in constant light (for condition B corresponding to one day after placing the cover, see Fig 1).

Rhythmicity and period length of activity, sleep, and position were analyzed using the Lomb-Scargle (LS) algorithm, with the same parameters to allow comparability (period range: 16 to 40 hours, resample rate: 30 minutes, alpha > 0.01). We visually compared periodograms and actograms to confirm rhythmicity. Period length and rhythmic strength were quantified using flies for which the LS algorithm identified a significant peak. Rhythmic strength of each fly was quantified as the difference between the power and the significance threshold. To generalize the observed differences between conditions in rhythmic strength, flies were further categorized in weakly or strongly rhythmic. To do so, flies that exhibited a rhythmic behavior with a relative power > 0.1 were considered strongly rhythmic while flies with a rhythmic strength < 0.1 were considered weakly rhythmic. This cut-off was selected because below 0.1 some flies in constant light without the cover (condition A) were considered rhythmic by the LS algorithm (Tab 1). Unless specified in the figure, we quantified weakly rhythmic flies together with arrhythmic flies. Sleep was defined as period of inactivity longer than 5 minutes *(30)*. Data were analyzed and plotted using Rethomics, a Rstudio package developed for the analysis of circadian behavior and sleep *(31)*. Flies that died before day 8 in constant light - or before the end of the experiment in the case of flies in DD - were excluded from the analysis.

.

*Immunohistochemistry*

Flies were loaded into the behaviour tubes in the Ethoscopes and entrained to 12hr:12hr LD for one day. Immediately after the transition to LL, the cover was placed (condition C). After 4 days, the behavior was analyzed, and the flies were classified as arhythmic or rhythmic as described above, and the activity status was estimated to predict center of the active and inactive phase on the next day. On the following day, flies of the two groups were then fixed in 4% PFA for 2.5 h at room temperature (RT). After fixation, the samples were washed 6 times with 0.1 M phosphate buffer (pH 7.4) with 0.1% Triton X-100 (PBS-T) at RT. Brains were dissected in PBS, were then blocked with 5% goat serum in 0.1% PBS-T for 2 h at RT and stained with pre-absorbed rabbit anti-PER (1:1000) *(32)* and mouse anti-PDF C7 (DSHB, 1:200) in 5% goat serum and 0.5% PBST for at least 48 h at 4ºC. After washing 3 times in PBS-T, the samples were incubated at 4°C overnight with goat anti-rabbit AlexaFluor 488 nm (1:500) and anti-mouse AlexaFluor 647 nm (1:500 Molecular Probes) in PBS-T. Brains were washed 3 times in PBS-T before being mounted in Vectashield. The images were taken using a Leica SP8 confocal microscope, keeping the same conditions for each experiment. For quantification, pixel intensity of mean and background staining in each neuronal group was measured by FIJI *(33)*. For each cell, three measurements were taken, as well as 3 measurements of the background for the corresponding section. Except for the DN3, intensity values were corrected for the number of total neurons stained, normalized by the known number of neurons of each group. Images were processed in GIMP.

*Statistics.*

Statistics analysis of behavioral data was conducted using Rstudio *(34)*. The statistical tests used are specified in each figure. We used t-test (paired and independent) for two- groups comparisons and ANOVA for multiple comparisons. Due to the differences in variance between compared groups determined by the Levene's test of Homogeneity of Variance, both ANOVA and t-test (paired and independent) were run with Welch correction. ANOVA was followed by the Games-Howell Post-Hoc test. Percentages and generalized linear model (GLM, family = binomial) was used to calculate whether there was an effect of condition on the number of rhythmic and arrhythmic flies, both for activity and position. Correlations were calculated using the Spearman's rank correlation test. Statistics are summarized in the supplementary tables 2-5). For comparison of staining intensity between active and inactive flies we applied estimation statistics *(16)*.

*Data and code availability*

Due to the size of the dataset, data will be provided by the corresponding author upon reasonable request. Data are stored on our local cloud server. R codes will be provided upon request

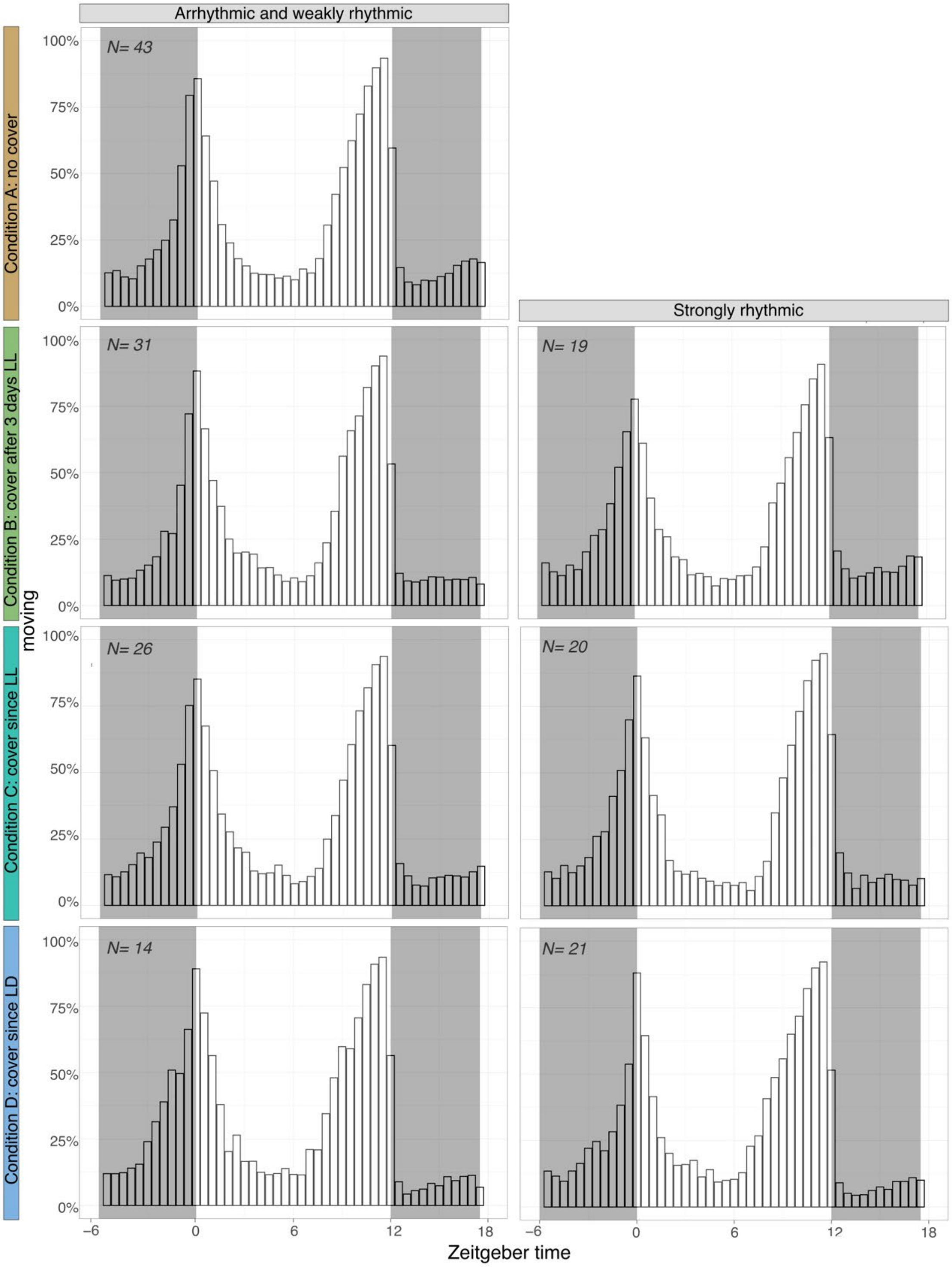


**Fig. S1. Behaviour of LL-arrhythmic and LL-rhythmic during the initial LD entrainment**. Daily average histograms of population activity of arrhythmic/weakly rhythmic flies and strongly rhythmic flies in each condition during initial LD entrainment (3 days). Light and dark phase are indicated by the white and the grey rectangles, respectively.

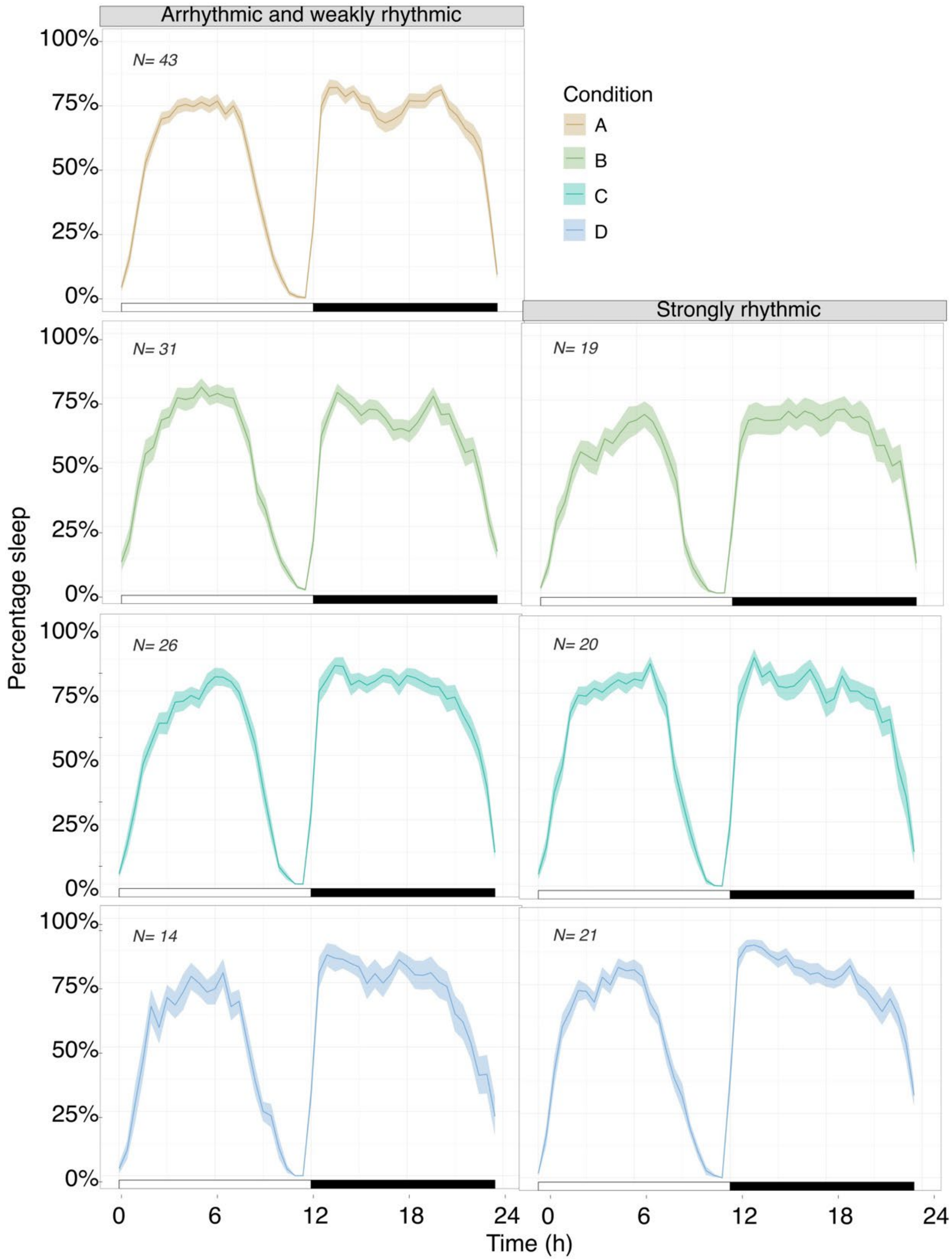


**Fig. S2. Normal sleep pattern of LL-arrhythmic and LL-rhythmic during the initial LD entrainment.** Daily average of population sleep of arrhythmic/weakly rhythmic flies and strongly rhythmic flies in each condition during the initial LD entrainment (3 days). Light and dark phase are indicated by the white and the black bar under each graph, respectively.
Create a page break and paste in the figure above the caption.

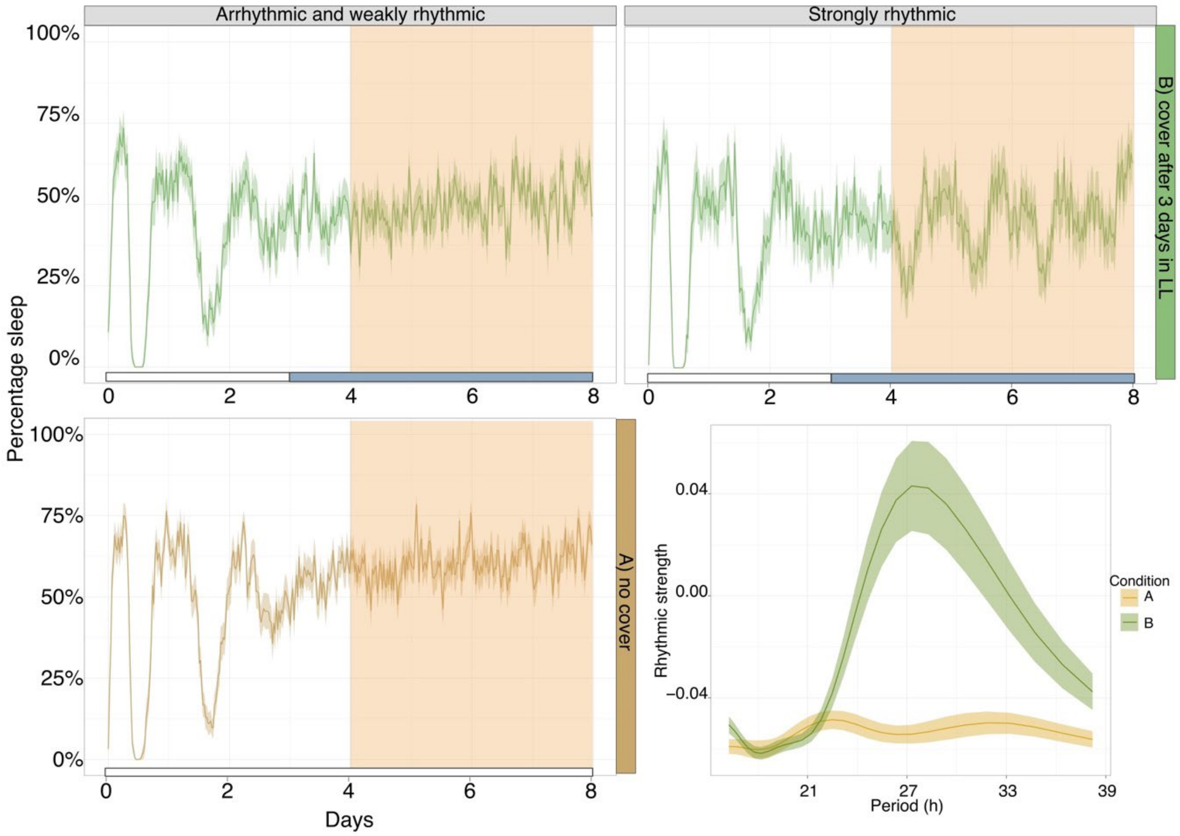


**Fig. S3. Sleep patterns of flies with arrhythmic locomotor activity compared to flies with restored rhythms in LL.** Percentage of sleep in of arrhythmic/weakly rhythmic (*top left; N= 31*) versus strongly rhythmic flies in condition B (*top right; N= 19*) and condition A (*bottom left; N= 43*). The sleep graphs from condition B are replotted from *Fig 5A*, to allow comparison with condition A. White and blue bars indicate the days in LL without or with cover. Lower right: Population periodogram comparing sleep rhythmic strength of condition A (*N= 43*) and B (*N= 50*).

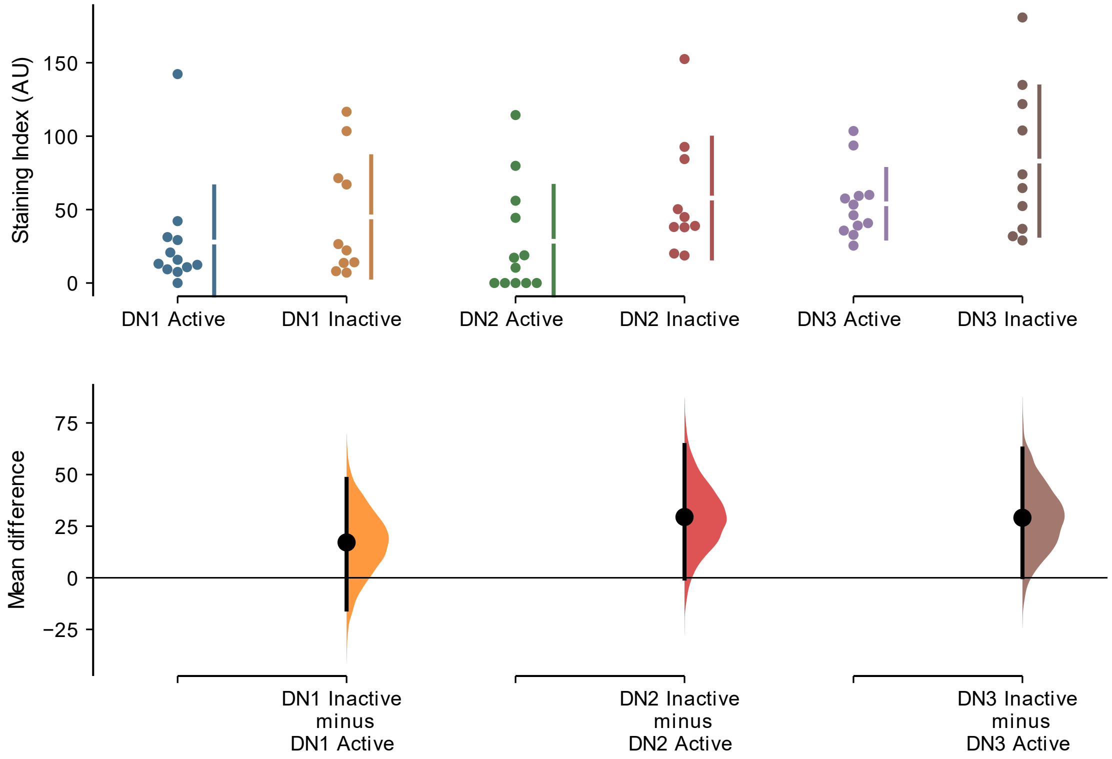


**Fig. S4. Quantification of PER expression in the dorsal clock neurons.** Rhythmic flies in LL (condition C) were dissected when they are either in their active or the inactive phase. The upper plot represents all the datapoints, with the mean (gap) and the standard deviation (length of vertical bars) for each neuronal group to the right. The lower plot shows the mean difference of the data from the inactive phase compared to the active phase as a bootstrap 95% confidence interval (see results text for details). n number: active phase 12, inactive phase 10, from 3 independent runs

| *iso31 flies in each condition after 4 days in constant light (LL)* | | | | | |
|---|---|---|---|---|---|
| Condition | RS | Total sleep (% ± SEM) | # sleep bouts (mean ± SEM) | Duration sleep bouts (secs) (mean± SEM) | Duration longest sleep bouts (secs) (mean± SEM) |
| No cover (A) | AR/WR | 63.4 ± 0.02 | 169.6 ± 6.7 | 1439.5 ± 104.2 | 10016.5 ± 1016.1 |
| Cover after 3 days LL (B) | AR/WR | 51.9 ± 0.03 | 178.9 ± 5.8 | 1017.1 ± 61.0 | 6551.6 ± 561.7 |
| | SR | 50.1 ± 0.04 | 172.2 ± 9.4 | 1120.3 ± 152.4 | 8464.2 ± 1406.2 |
| Cover since LL (C) | AR/WR | 61.4 ± 0.02 | 165.3 ± 6.2 | 1339.4 ± 79.5 | 8713.5 ± 851.7 |
| | SR | 59.8 ± 0.02 | 143.4 ± 6.8 | 1536.9 ± 124.4 | 13639.5± 1539.3 |
| Cover since LD (D) | AR/WR | 64.7 ± 0.03 | 155 ± 7.8 | 1506.4 ± 121.3 | 10954.3 ± 2211.5 |
| | SR | 65.0 ± 0.01 | 127.7 ± 4.3 | 1802.947 ± 69.8 | 16724.8 ± 1561.6 |

*Notes: AR=* Arrhythmic flies*; SR:* flies with a strong rhythm; *WR:* flies with a weak rhythm.

**Table S1. Sleep quality of strongly rhythmic vs. arrhythmic/weakly rhythmic wild type flies (iso31) in constant light with and without the cover.**

| *Parameters* | *Group 1 vs. group 2* | *Df1* | *Df2* | *Statistic* | *p-value* | *Significance* | *Fig.* |
|---|---|---|---|---|---|---|---|
| RS.act | | 3 | 97 | 7.77 | < .001 | *** | 2C |
| period.act | | | | 6.72 | < .001 | *** | 2D |
| RS.pos | | 3 | 137 | 6.26 | < .001 | *** | 3C |
| period | Activity vs position | 1 | 234 | 16.16 | < 0.0001 | **** | 4A-B |
| RS | | | | 17.32 | < 0.0001 | **** | 4C-D |
| n_bouts | B_AR/WR vs. B_SR | 1 | 48 | 1.25 | 0.27 | ns | 5B |
| | C_AR/WR vs. C_SR | 1 | 44 | 0.39 | 0.54 | ns | |
| | D_AR/WR vs. D_SR | 1 | 33 | 5.16 | 0.03 | * | |
| mean_bout | B_AR/WR vs. B_SR | 1 | 48 | 5.09 | 0.03 | * | 5C |
| | C_AR/WR vs. C_SR | 1 | 44 | 0.33 | 0.57 | ns | |
| | D_AR/WR vs. D_SR | 1 | 33 | 0.65 | 0.43 | ns | |
| longest_bout | B_AR/WR vs. B_SR | 1 | 48 | 3.90 | 0.05 | ns | 5D |
| | C_AR/WR vs. C_SR | 1 | 44 | 6.44 | 0.01 | * | |
| | D_AR/WR vs. D_SR | 1 | 33 | 0.08 | 0.79 | ns | |

*Abbreviations: RS.act:* rhythmic strength activity; *period.act:* period length activity; *RS.pos:* rhythmic strength positional change; *period.pos:* period length positional change; *AR/WR:* arrhythmic/weakly rhythmic flies; *SR:* strongly rhythmic flies; *n_bouts:* number of sleep bouts; *mean_bout:* average duration of one sleep bout; *longest_bout:* duration of the longest bout. **** $p < .0001$, *** $p < .001$, ** $p < .01$, * $p < .05$, ns $p > .05$.

**Table S2. Levene test for Homogeneity of Variance for each figure.**

| *Response variable tested* | *Coefficients* | *Deviance residuals* | | | | | *Estimate* | *SE* | *Z-value* | *p-value* | *Signif.* | *Fig.* |
|---|---|---|---|---|---|---|---|---|---|---|---|---|
| | | *Min* | *1Q* | *Median* | *3Q* | *Max* | | | | | | |
| rhythmic.act | | -1.72 | -0.81 | 0.72 | 0.89 | 1.60 | | | | | | 2B |
| | intercept | | | | | | -0.95 | 0.34 | -2.79 | 0.005 | ** | |
| | Condition B | | | | | | 1.44 | 0.45 | 3.212 | 0.001 | ** | |
| | Condition C | | | | | | 1.67 | 0.46 | 3.62 | 0.0003 | *** | |
| | Condition D | | | | | | 2.16 | 0.53 | 4.11 | < 0.0001 | **** | |
| rhythmic.pos | | -2.06 | 0.51 | 0.57 | 0.67 | 0.85 | | | | | | 3B |
| | intercept | | | | | | 0.84 | 0.33 | 2.52 | 0.01 | * | |
| | Condition B | | | | | | 1.16 | 0.55 | 2.11 | 0.03 | * | |
| | Condition C | | | | | | 0.880 | 0.53 | 1.67 | 0.09 | ns | |
| | Condition D | | | | | | 0.55 | 0.54 | 1.02 | 0.31 | ns | |

*Abbreviations: rhythmic.act:* flies that shows rhythmic activity; *rhythmic.pos:* flies that shows a rhythmic change in position within the tube. **** $p < .0001$, *** $p < .001$, ** $p < .01$, * $p < .05$, ns $p > .05$.

**Table S3. Summaries for Generalised Linear model (GLM), with deviance residuals.**

| Parameter | Group1 vs. group 2 | N. n1 | N. n2 | Test | Estimates | *p-value* | *Signif.* | *Fig.* |
|---|---|---|---|---|---|---|---|---|
| RS.act | - | 101 | | ANOVA | $F_{(3,51.51)}$=38.12 | < .0001 | **** | 2C |
| period.act | - | 101 | | | $F_{(3,37.03)}$= 2.51 | 0.07 | ns | 2D |
| RS.pos | - | 141 | | | $F_{(3,68.37)}$= 13.08 | < .0001 | **** | 3C |
| Condition B | period.act vs period.pos | 38 | | Paired t-test | *t(37)*= -1.83 | 0.23 | ns | 4A |
| Condition C | period.act vs period.pos | 40 | | | *t(39)*= 2.85 | 0.02 | * | |
| Condition D | period.act vs period.pos | 40 | | | *t(39)*= 1.11 | 0.82 | ns | |
| Condition B | period.act vs period.pos | 38 | | Sperman's rank rho correlation test | *S*= 185.8, *ρ*= 0.837 | < 0.0001 | **** | 4B |
| Condition C | period.act vs period.pos | 40 | | | *S*= 894.3, *ρ*= 0.328 | 0.16 | ns | |
| Condition D | period.act vs period.pos | 40 | | | *S*= 803.5, *ρ*= 0.396 | 0.08 | ns | |
| Condition B | RS.act vs RS.pos | 38 | | Paired t-test | *t(37)*= -1.03 | 0.94 | ns | 4C |
| Condition C | RS.act vs RS.pos | 40 | | | *t(39)*= 1.64 | 0.32 | ns | |
| Condition D | RS.act vs RS.pos | 40 | | | *t(39)*= 2.67 | 0.03 | * | |
| Condition B | RS.act vs RS.pos | 38 | | Sperman's rank rho correlation test | *S*= 360, *ρ*= 0.684 | 0.01 | ** | 4D |
| Condition C | RS.act vs RS.pos | 40 | | | *S*= 640, *ρ*=0.519 | 0.02 | * | |
| Condition D | RS.act vs RS.pos | 40 | | | *S*= 286, *ρ*= 0.785 | < 0.0001 | **** | |
| n_bouts | B_weak vs B_strong | 19 | 31 | Independent t-test | *t(31.6)*= -0.61 | 0.55 | ns | 6B |
| | C_weak vs C_strong | 20 | 26 | | *t(41.7)*= -2.38 | 0.02 | * | |
| | D_weak vs D_strong | 21 | 14 | | *t(20.9)*= -3.07 | 0.01 | ** | |
| mean_bout | B_weak vs B_strong | 19 | 31 | | *t(23.9)*= 0.63 | 0.54 | ns | |
| | C_weak vs C_strong | 20 | 26 | | *t(33.4)*= 1.34 | 0.19 | ns | |
| | D_weak vs D_strong | 21 | 14 | | *t(21.5)*= 2.12 | 0.05 | * | |
| longest_bout | B_weak vs B_strong | 19 | 31 | | *t(23.8)*= 1.26 | 0.22 | ns | |
| | C_weak vs C_strong | 20 | 26 | | *t(30.3)*= 2.80 | 0.01 | ** | |
| | D_weak vs D_strong | 21 | 14 | | *t(25.1)*= 2.13 | 0.04 | * | |

*Note:* ANOVA, paired and independent t-test were corrected with Welch correction method. *Abbreviations: RS.act:* rhythmic strength activity; *period.act:* period length activity; *RS.pos:* rhythmic strength positional change; *period.pos:* period length positional change; *n_bouts:* number of sleep bouts; *mean_bout:* average duration of one sleep bout; *longest_bout:* duration of the longest bout. **** *p* < .0001, *** *p* < .001, ** *p* < .01, * *p* < .05, ns *p* >.05.

**Table S4. ANOVA, Paired t-test with Welch's correction and Spearman's Rank Sum rho correlation test**

| *Parameter* | *Group 1* | *Group 2* | *Mean ± SE* | *95% CI* | | *p-value* | *Significance* | *Fig* |
|---|---|---|---|---|---|---|---|---|
| | | | | *low* | *high* | | | |
| RS.act | A (N=12) | B (N=31) | 0.15±0.02 | 0.08 | 0.23 | < .0001 | **** | 2C |
| | | C (N= 31) | 0.18±0.02 | 0.11 | 0.26 | < .0001 | **** | |
| | | D (N= 27) | 0.22±0.02 | 0.15 | 0.30 | < .0001 | **** | |
| | B (N=31) | C (N= 31) | 0.03±0.03 | -0.07 | 0.14 | 0.84 | ns | |
| | | D (N= 27) | 0.07±0.03 | -0.03 | 0.18 | 0.26 | ns | |
| | C (N=31) | D (N= 27) | 0.04±0.03 | -0.07 | 0.15 | 0.76 | ns | |
| period.act (secs) | A (N=12) | B (N=31) | 4645.18±4670.25 | -14596.97 | 23887.34 | 0.89 | ns | 2D |
| | | C (N= 31) | 8407.59±4540.22 | -10579.87 | 27395.06 | 0.57 | ns | |
| | | D (N= 27) | 11428.00±4582.14 | -7638.44 | 30494.43 | 0.33 | ns | |
| | B (N=31) | C (N= 31) | 3762.41±1872.46 | -3258.37 | 10783.19 | 0.49 | ns | |
| | | D (N= 27) | 6782.81±1971.92 | -605.72 | 14171.34 | 0.08 | ns | |
| | C (N=31) | D (N= 27) | 3020.40±1640.37 | -3130.83 | 9171.63 | 0.57 | ns | |
| RS.pos | A (N=30) | B (N=44) | 0.11±0.02 | 0.04 | 0.18 | < .001 | *** | 3C |
| | | C (N= 39) | 0.11±0.02 | 0.03 | 0.18 | < .01 | ** | |
| | | D (N= 28) | 0.15±0-02 | 0.06 | 0.24 | < .001 | *** | |
| | B (N=27) | C (N= 39) | -0.01±0.03 | -0.10 | 0.09 | 1.00 | ns | |
| | | D (N= 28) | 0.04±0.03 | -0.07 | 0.14 | 0.81 | ns | |
| | C (N=39) | D (N= 28) | 0.04±0.03 | -0.07 | 0.15 | 0.73 | ns | |

*Abbreviations: RS.act:* rhythmic strength activity; *period.act:* period length activity; *RS.pos:* rhythmic strength positional change.
**** $p < .0001$, *** $p < .001$, ** $p < .01$, * $p < .05$, ns $p > .05$.

**Table S5. ANOVA, Paired t-test with Welch's correction and Spearman's Rank Sum rho correlation test**

| *Neuronal group* | *Phase* | *Mean±SEM* | *Difference to the active phase* | *95% CI* | |
|---|---|---|---|---|---|
| | | | | low | *high* |
| lLNv | Active | 40.56 ± 6.9 | | | |
| | Inactive | 69.99 ± 14.4 | 29.4 | 1.27 | 59.8 |
| sLNv | Active | 41.92 ± 7.8 | | | |
| | Inactive | 67.15 ± 13.4 | 25.2 | -0.17 | 56.7 |
| 5$^{th}$ sLNv | Active | 52.44 ± 13.9 | | | |
| | Inactive | 70.46 ± 19.9 | 18.0 | -28.2 | 62.3 |
| LNd | Active | 57.63 ± 10.0 | | | |
| | Inactive | 103.84 ± 17.6 | 46.2 | 10.6 | 83.7 |
| DN1 | Active | 27.89 ± 10.9 | | | |
| | Inactive | 45.02 ± 13.1 | 17.1 | -15.4 | 47.9 |
| DN2 | Active | 28.43 ± 10.9 | | | |
| | Inactive | 57.86 ± 13.0 | 29.4 | -0.37 | 64.3 |
| DN3 | Active | 53.95 ± 6.8 | | | |
| | Inactive | 83.02 ± 16.1 | 29.1 | 0.32 | 62.6 |

**Table S6. Unpaired mean differences calculated by estimation statistics**
